# Revisiting Mouse Cardiac Myocyte Isolation: A Simplified Langendorff-based Method

**DOI:** 10.64898/2026.04.15.718810

**Authors:** Mie Sarah Larsen, Morten B Thomsen, Tamzin Zawadzki

## Abstract

This protocol describes a Langendorff-based method for isolating intact adult mouse ventricular myocytes using syringe pump-driven perfusion. The approach retains the key physiological advantage of the conventional Langendorff technique, continuous retrograde coronary perfusion, while simplifying the overall setup. By combining retrograde aortic perfusion with widely available laboratory equipment, the method provides an accessible alternative to traditional Langendorff systems. A precision syringe pump connected to an in-line heater is used to deliver temperature-controlled, constant-flow perfusion during enzymatic digestion. In contrast to gravity-driven constant-pressure systems, constant-flow perfusion maintains stable enzyme delivery despite changes in coronary resistance that occur during tissue digestion. Use of an inline heater allows precise, rapid temperature-controlled delivery, avoiding the complexity, leak risk, thermal lag, and contamination susceptibility associated with traditional water-jacketed systems. Our setup reduces variability in perfusion rate and minimizes susceptibility to occlusion, flow interruption, or compliance-related artifacts, enhancing reproducibility. The method consistently yields adult ventricular myocytes with high viability (>70% rod-shaped, calcium-tolerant), enabling a broad range of functional analyses including electrophysiology, contractile performance and calcium handling. Step-by-step instructions, troubleshooting guidance, and anticipated outcomes are provided to facilitate adoption in laboratories without dedicated isolated-heart perfusion infrastructure.

**Key Features:**

- Simplified Langendorff-based mouse cardiomyocyte isolation method that eliminates the need for specialized perfusion rigs.
- Syringe pump–driven constant-flow perfusion combined with inline temperature control improves reproducibility by ensuring stable enzyme delivery and precise temperature regulation.
- Generates high-yield, calcium-tolerant adult mouse ventricular myocytes suitable for functional studies.

**Graphical Overview:** Graphical overview of the simplified Langendorff-based mouse cardiac myocyte isolation protocol.

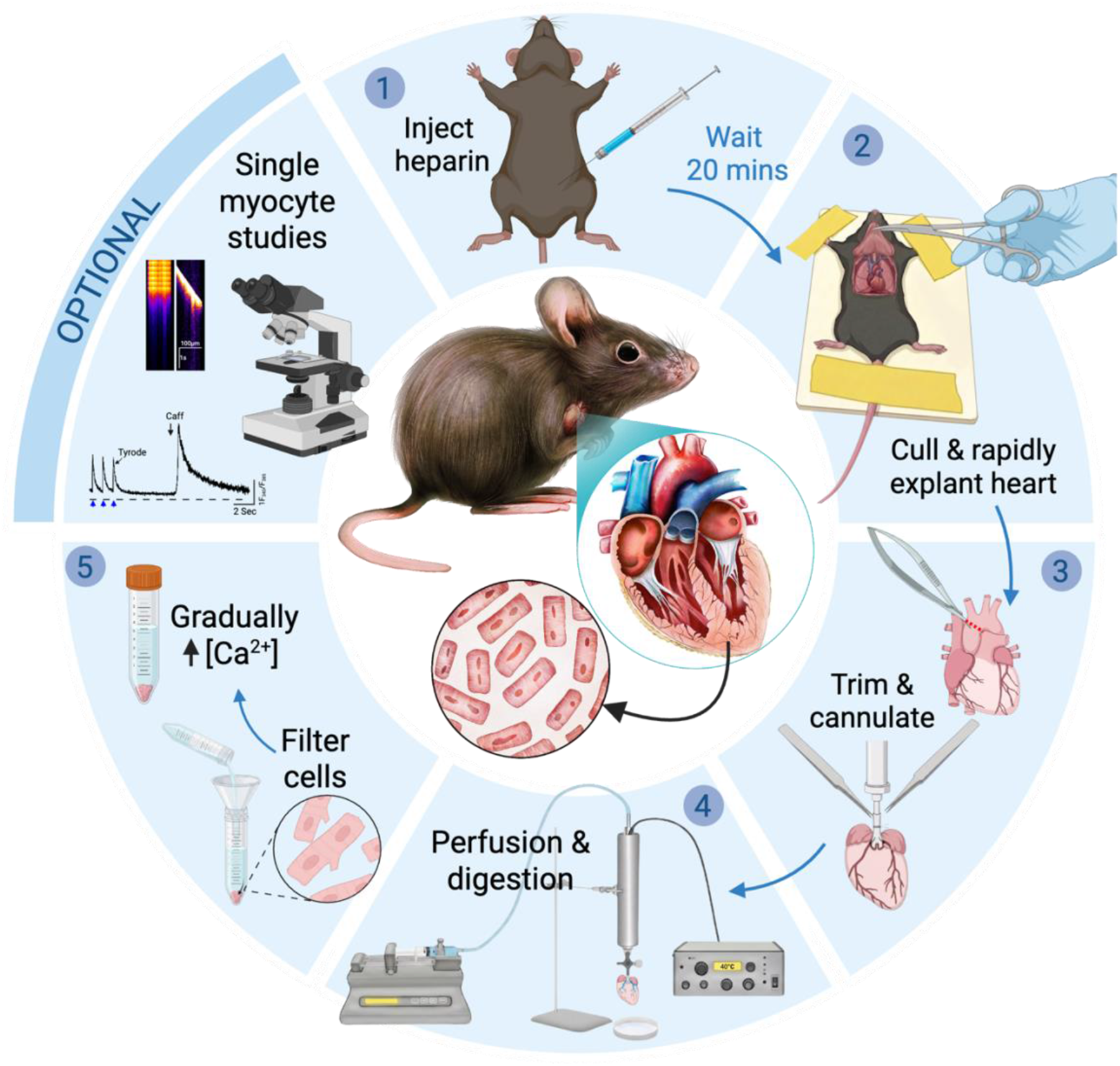

## Background

The isolation of intact adult cardiac myocytes is a foundational technique in cardiac research, enabling direct investigation of cellular structure and function under controlled experimental conditions. The isolated heart preparation described by Oscar Langendorff in 1897,^1^ and methodologically refined by others,^2,3^ forms the basis of the modern “Langendorff” method.^4^ In this preparation, the excised heart is rapidly cannulated via the ascending aorta and perfused *ex vivo* while suspended in a temperature-controlled chamber. Retrograde perfusion through the aorta allows controlled delivery of physiological solutions through the intact coronary vasculature while preserving myocardial architecture.^1^ Its further adaptation in the 1970s for enzymatic dissociation established the Langendorff method as the standard approach for isolating viable cardiac myocytes,^5,6^ a status it has maintained for more than five decades.^7–9^ Langendorff perfusion enables enzymatic delivery through the coronary circulation while minimizing mechanical trauma to the myocardium by retrograde aortic perfusion. In this configuration, flow delivered through the cannulated proximal aorta fills the aortic root, generating pressure that causes coaptation of the aortic cusps and closure of the valve. Aortic valve closure prevents retrograde ventricular filling and directs perfusate into the coronary circulation via the coronary ostia at the base of the root.^10^ This technique is particularly well suited to rodents and other small-animal models, in which the aorta is sufficiently large for reliable cannulation yet small enough to allow economical use of dissociation enzymes.^11^ For larger hearts, perfusing a coronary branch and excising the region is more practical than perfusing the whole heart.^12^

Langendorff systems operate under either constant-pressure or constant-flow perfusion regimes.^8^ Constant-pressure setups are typically gravity-fed from a reservoir of perfusate and exhibit variable coronary flow as vascular resistance changes during digestion. As enzymatic dissociation reduces resistance, coronary flow increases, providing a useful indicator of digestion-progression that is commonly monitored by changes in drip rate.^11,13–16^ Because coronary vascular resistance dynamically influences the pressure–flow relationship,^17^ variations in flow under constant-pressure perfusion may result in variable delivery of enzyme, particularly as resistance drops during digestion.^11,16^ Variability in extracellular matrix composition and fibrosis degree across strain, age, and pathological conditions ostensibly will also influence myocardial resistance and thereby perfusion flow dynamics.^16^

In contrast, constant-flow systems employ a fixed volumetric perfusion rate while perfusion pressure varies in response to downstream resistance. Careful selection of the flow rate is important: if set too low, inadequate coronary perfusion may occur; if excessive, supraphysiological perfusion pressures can invert the aortic valve cusps and divert perfusate into the ventricular cavity rather than the coronary circulation. Many traditional constant-flow Langendorff setups rely on peristaltic pumps as well as dedicated perfusion rigs incorporating water-jacketed glassware, bubble traps, recirculation loops, and continuous carbogen or oxygen gassing of bicarbonate-buffered solutions.^5,6,9,13^ The complexity of specialized apparatus, combined with the technical expertise required for rapid aortic cannulation, has been criticized as limiting accessibility. In response, simplified “Langendorff-free” methods have been developed.^18–21^

Langendorff-free methods, sometimes referred to as ‘injection methods’, are based on direct injection of perfusate into the ventricular lumen.^19–22^ In these approaches, the aorta is clamped, and buffer is forced into the coronary circulation in an antegrade manner, flowing through the aortic valve into the coronary ostia. Although injection–based methods are touted as requiring less technical skill than aortic cannulation, they nonetheless depend on accurate clamp placement and precise ventricular injections, first into the right ventricular lumen followed by multiple injections into the left ventricular lumen. These procedures demand operator precision and introduce distinct risks. Repeated ventricular punctures, particularly in the small and thin-walled mouse heart, introduce the risk of rupture or buffer leakage. Preservation of aortic integrity is also critical, as damage can compromise effective coronary perfusion. These risks may be exacerbated in structurally fragile or diseased hearts or those exhibiting pathological remodeling.

Moreover, injection methods rely on non-physiological alkaline buffers (pH ∼7.8) for optimal cell yield, ^18–20^ the basis for which is speculative rather than mechanistically established.^19^ Cardiomyocyte function is sensitive to pH, and extracellular alkalinization directly influences intracellular pH via established acid–base transport mechanisms, including Na^+^/H^+^ exchange and lactate/H^+^ transport.^23,24^ Because intracellular pH must be tightly maintained within a narrow physiological range (∼7.15–7.25) to preserve excitation–contraction coupling and calcium homeostasis, even modest deviations can alter Ca^2+^ handling and myofilament sensitivity.^25–28^ Furthermore, extracellular pH fluctuations regulate autophagy, a process central to cellular viability and metabolic control.^29^ Exposure to alkaline conditions during isolation may influence post-isolation cellular physiology and subsequent functional readouts.

Taken together, these considerations highlight the need for a method that preserves the physiological fidelity and controlled delivery afforded by constant-flow coronary perfusion while eliminating the specialized equipment and technical expertise required by traditional Langendorff systems. The protocol described here addresses this need through a simplified, syringe pump–driven Langendorff approach. It retains the critical advantages of aortic cannulation and continuous retrograde perfusion while implementing them within a minimal and accessible apparatus optimized for the isolation of adult mouse ventricular myocytes.

## Materials

### Reagents & Materials

| Item | Example Supplier | Catalog # | Notes |
| --- | --- | --- | --- |
| 25, 100 mL beaker | Corning/PYREX® | 1395-25<br>1395-100 |  |
| 50ml volumetric flask | Corning/PYREX® | CLS564050FO |  |
| Black braided silk<br>USP size 5-0 | Galerie Vömel GmbH | 14739 |  |
| Cell strainer | pluriStrainer® 300µm,<br>pluriSelect Life Science | 43-50300-03 |  |
| Analytical funnel | Semadeni | 224 | Diameter ø 37 mm |
| Conical screw cap<br>tube, 15 ml | Sarstedt | 62.554.502 |  |
| Pasteur pipette, 3 ml | Auxilab | DLB009 |  |
| 1ml syringe | BD Plastipak Luer-lock<br>precision syringe |  |  |
| Luer-lock needle | 26G x 1” | 4710004525 |  |
| 10ml syringe | B.Braun | 4617100V-02 |  |
| Collagenase II | Worthington Biochemical | LS004177 | Target 6500 U per 50 mL;<br>store 4 °C |
| Collagenase IV | Worthington Biochemical | LS004189 | Target 5250 U per 50 mL;<br>store 4 °C |
| Protease XIV | Merck | P5147 | 2.5 mg per 50 mL; store - 20 °C |
| CaCl<sub>2</sub> solution (1 M stock) | Merck | 21115 | Dilute to 10 mM and 100 mM working stocks |

### Equipment

| Equipment | Example Model/Brand | Notes |
| --- | --- | --- |
| Analytical balance | Mettler AT261 DeltaRange®/ Mettler Toledo |  |
| Magnetic stirrer | IKA-COMBIMAG RCO/ IKA |  |
| Water bath | Techne Tempette TE-8A Water Bath |  |
| Syringe pump | KD Scientific Model 100 | Controlled perfusion |
| Inline heater and temperature controller | Warner Instruments: SH-27B inline heater and TC-324B | Set to 40°C; adjust so delivers 37°C at needle tip |
| Dissecting microscope | ZEISS Stemi 305 | Used for cannulation |
| Cannula 23G/ 25G | Custom notches/ridges to secure the suture during tying | Prevents slippage and stabilizes the cannulated heart |
| Tissue forceps (rat-tooth forceps) | Aesculap #BD557R | 1×2 teeth, 145mm |
| Iris scissors | Aesculap #BC110R |  |
| Spring micro scissor | Vannas-Tübingen Spring Scissors# 15003-08 | Kept protected in silastic tube or pipette tip |
| Operating scissor | KRUUSE 14cm #130460 |  |
| Hemostatic forceps, curved | Aesculap #BH111R | Also called Mosquito forceps |
| Curved iris forceps, curved 4” serrated tip | Graefe, CHIRU+, Surgical Instruments #19-7013.01 |  |
| No. 5 Fine Forceps (Stainless Steel) x 2 | Dumont, Merck #F6521 | Smooth tips, for delicate tissue. Kept protected in silastic tube |

### Recipes

Ca^2+^-free Tyrode’s solution (136 mM NaCl, 4 mM KCl, 5 mM HEPES, 5 mM MES, 0.8 mM MgCl_2_, 10 mM glucose; 37 °C; pH = 7.4). Minimum 10ml.

EDTA buffer (125 mM NaCl, 5 mM KCl, 0.5 mM NaH_2_PO_4_, 10 mM HEPES, 10 mM BDM, 10 mM taurine, 5 mM EDTA, 10 mM glucose; 37 °C; pH = 7.4). Minimum 25ml.

Perfusion buffer (130 mM NaCl, 5 mM KCl, 0.5 mM NaH_2_PO_4_, 10 mM HEPES, 10 mM BDM, 10 mM taurine, 1 mM MgCl_2_, 10 mM glucose; 37 °C; pH = 7.4). Minimum 50ml.

Enzyme solution (perfusion buffer supplemented with collagenase II [3250 U per 25 mL], collagenase IV [2625 U per 25 mL], and protease XIV [1.25 mg per 25 mL]; 37 °C; pH = 7.4). Minimum 25ml.

Stop buffer (perfusion buffer supplemented with 5% w/v BSA; 37 °C; pH = 7.4). Minimum 10ml.

Chemicals from Sigma-Aldrich.

### Ethical Approval

All procedures described in this protocol were approved by the Danish National Animal Experiments Inspectorate under the Danish Ministry of Environment and Food (license #2023-15-0201-01576) and conducted in accordance with European Parliament Directive 2010/63/EU on the protection of animals used for scientific purposes. All experiments complied with applicable institutional and national animal welfare regulations. Mice were euthanized by cervical dislocation performed by trained personnel in accordance with the approved project license. Investigators implementing this protocol must obtain appropriate ethical approval and ensure that all animal procedures, including euthanasia methods, comply with relevant institutional and national regulations prior to initiating experiments.

### Procedure

1. Inject anticoagulant heparin sodium at 1000 U/mL, 0.2 mL intraperitoneally (200 U per mouse; ∼10 U/g for a 20 g mouse) 20 min prior to euthanasia.
2. Euthanize mouse by cervical dislocation. Quickly tape front limbs and tail to immobilize in supine position. From the moment of respiratory arrest oxygen delivery to the heart is interrupted. Within 10-30 seconds anoxia and ischemia are followed by dramatic changes in ATP and creatine phosphate content.^30^ The following steps should therefore be conducted rapidly until consistent coronary perfusion is achieved (step 9).
3. Lift skin at sternum with tissue (rat tooth) forceps and cut using blunt-end operating scissors. Reflect skin caudally and cranially to expose abdominal cavity.
4. Make incision under ribcage, cutting diaphragm; bilaterally cut to retroflect thoracic cage with hemostatic forceps, exposing heart.
5. Elevate the heart gently with curved iris forceps and excise, keeping dissection with iris scissors close to the dorsal thoracic wall to preserve aortic length. Sever the inferior vena cava and descending aorta first to facilitate rapid mobilization of the heart while minimizing tension on the ascending aorta. Remove the lungs, thymus, and pericardial fat pads, ensuring the heart, including the ascending aorta and atria, remains intact.
6. Transfer heart to ice-cold EDTA buffer in cannulation chamber under dissecting microscope. Trim using spring micro scissors (Figure 1). There is no need to ‘rinse’ the heart to remove blood as this unnecessarily extends the period of ischemia. In sufficiently heparinized animals, latent blood will be washed out after the onset of perfusion. Any thrombi formed in the arterial coronary vasculature will not be removed by rinsing anyway.
7. Insert a 25G cannula into the aorta and gently advance it towards the aortic root, positioning the tip just above the aortic valve. Using fine forceps, guide the aorta over the cannula. The cannula and perfusion line should be pre-filled with EDTA buffer and free of air bubbles. Secure the aorta in place with a double suture (reef knot).
8. Transfer cannulated heart to syringe pump (e.g., KD Scientific Model 100) capable of delivering stable, low flow rates. See Figure 3.
9. Perfuse EDTA buffer at 1 mL/min for 5 min. Meanwhile, dissolve enzyme aliquots to prepare Enzyme solution
10. Switch to Perfusion buffer at 1.5 mL/min for 2 min.
11. Perfuse Enzyme solution at 2 mL/min for ∼5 min. Monitor digestion progress continuously. The heart should become progressively swollen, softer, and flaccid. A slight shift from a reddish/dark pink to pale pink is expected during proper digestion. As digestion proceeds, the tissue should feel soft and spongy when gently pinched with forceps, and the surface may show loosening or separation of muscle fibers. *Note:*Pale pink colouration during digestion should not be confused with a white or yellowish appearance, which may indicate poor coronary perfusion (e.g., incorrect cannulation or air embolism).
12. Endpoint: Stop perfusion when the heart is clearly soft, flaccid, and spongy, while maintaining a light pink (not white) appearance. Transfer the heart to a new dish containing ∼5 mL stop buffer (perfusion buffer + 5% BSA).
13. Gently tease tissue with forceps. Well-digested tissue is pale, soft, and tears easily between forceps to release cells. If tissue appears firm and dark, this indicates ineffective coronary perfusion, insufficient enzymatic digestion, or both. Optionally use scissors in feathering motion and cut-tip Pasteur pipette for gentle trituration (avoid bubbles as shear stress can damage cell membranes).
14. Filter suspension through 300 µm mesh into 15ml conical tube. Allow cells to pellet by gravity (∼12 min) or by centrifugation for 1 minute at 300RPM. Aspirate supernatant to discard and resuspend cell pellet in Ca^2+^-free Tyrode’s at RT.
15. Assess cell viability under microscope (10×). Acceptable if >70% rod-shaped cells. NB: To prevent sample contamination, microbial growth, salt precipitation, and flow obstruction, all tubing must be carefully maintained. After each use, tubing should be flushed with 70% ethanol followed by Milli-Q or ultra-pure water. Tubing should be inspected regularly and replaced at defined intervals to ensure consistent flow performance and experimental reproducibility. Tubing should be replaced immediately if discoloration, increased back pressure, reduced flow, or visible residue is observed.

**Figure 1.**
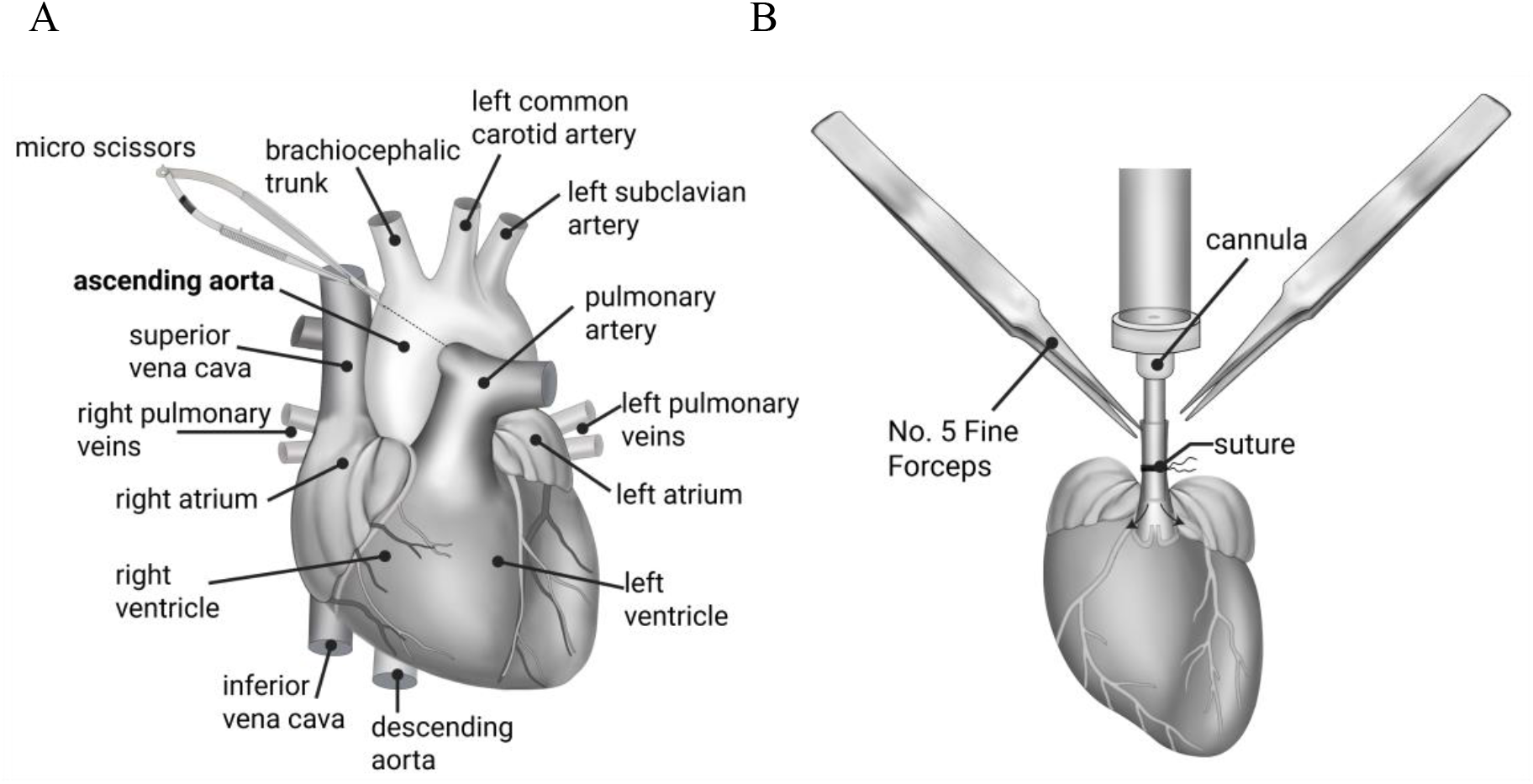
Preparation to cannulate aorta (A) Ascending aorta is trimmed using micro scissors beneath the brachiocephalic trunk to preserve a sufficient length of aorta for cannulation. (B) aorta is guided onto cannula using fine forceps and fixed in place with suture.

**Figure 2.**
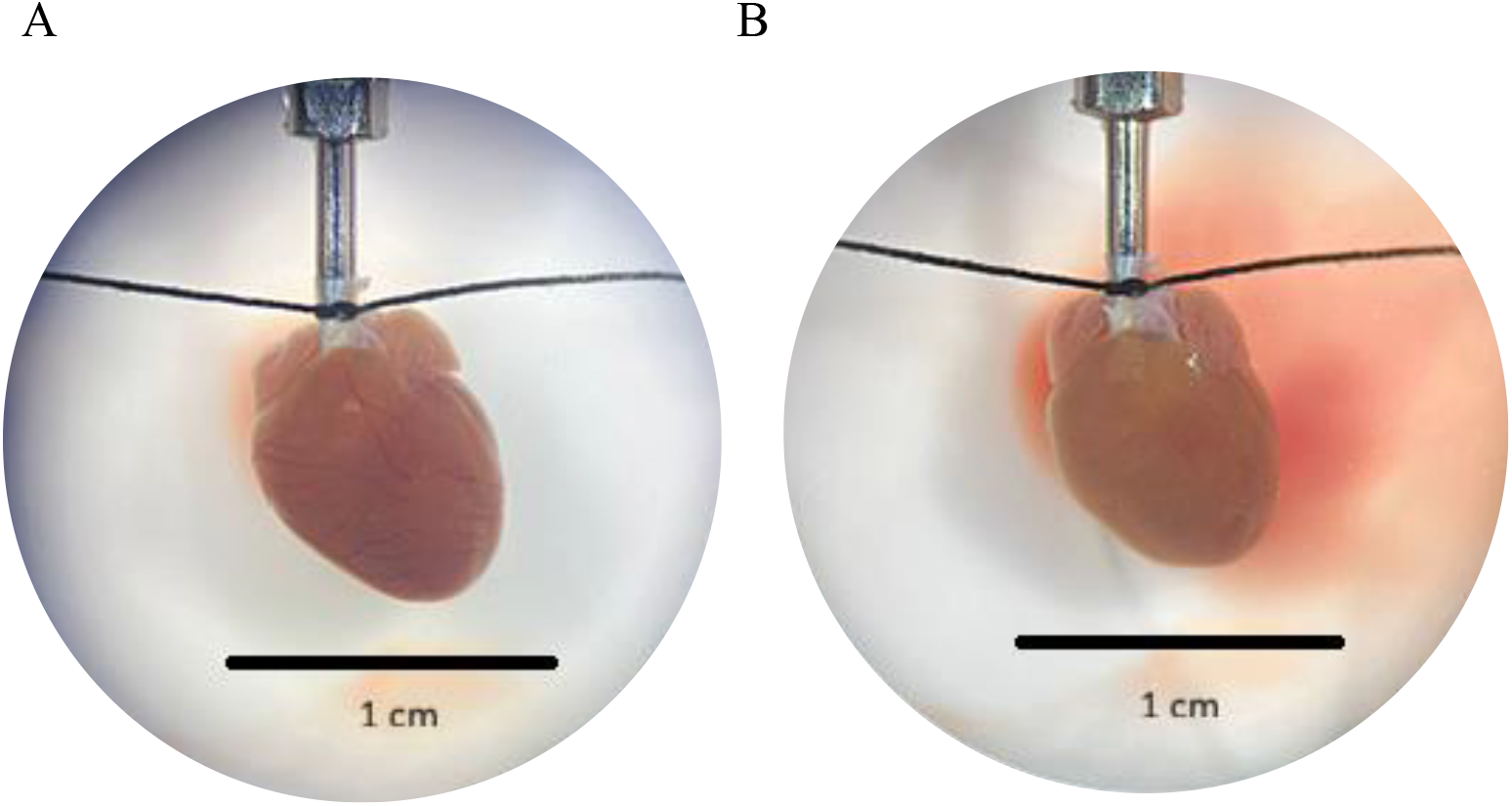
Mouse heart cannulation under microscope (10X). A) The aorta was secured on the cannula using silk suture thread. B) after flushing with EDTA buffer the coronary veins are cleared of blood.

**Figure 3.**
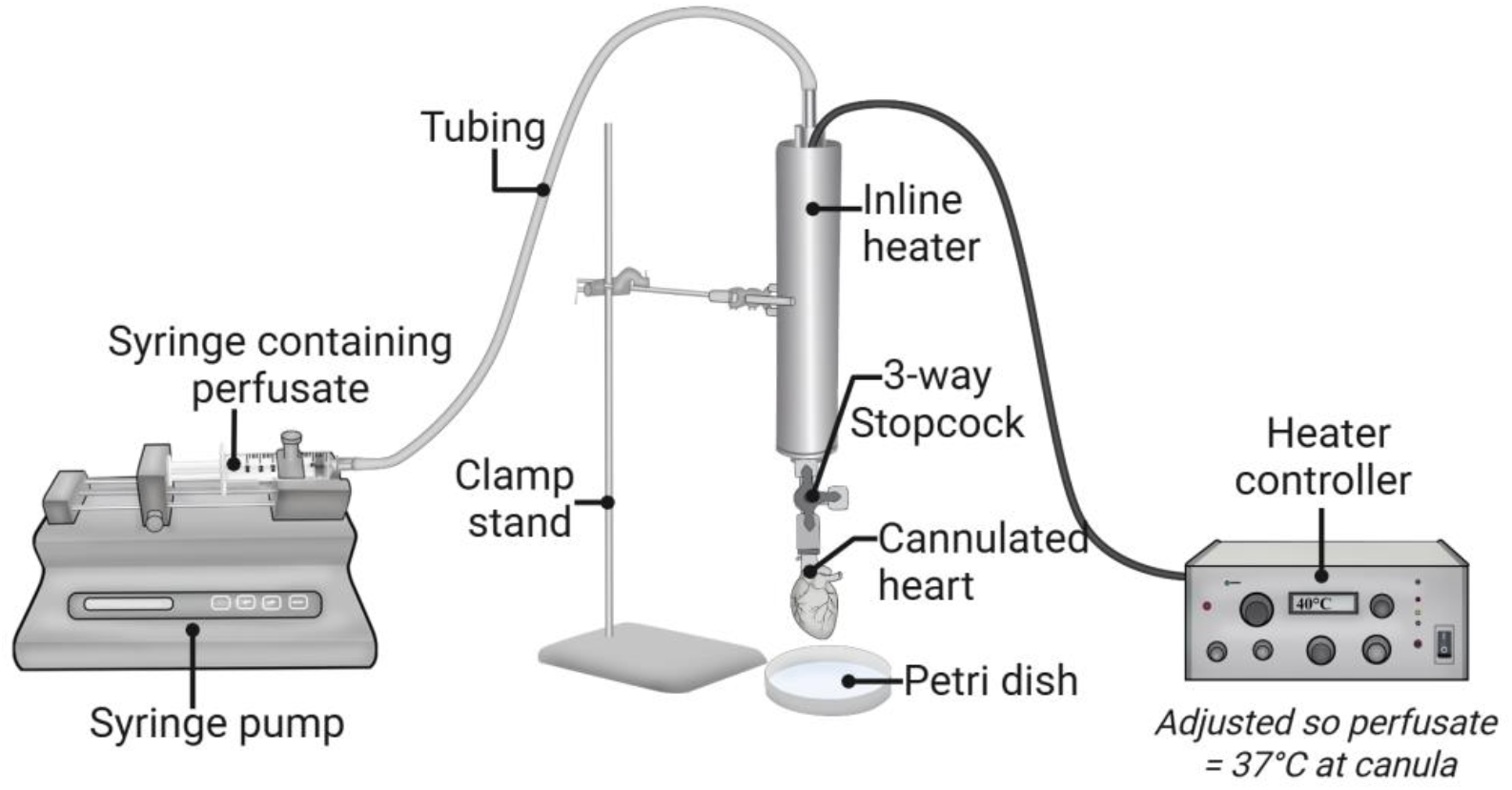
Cannulated heart is transferred to setup with syringe-pump and inline heater.

**Figure 4.**
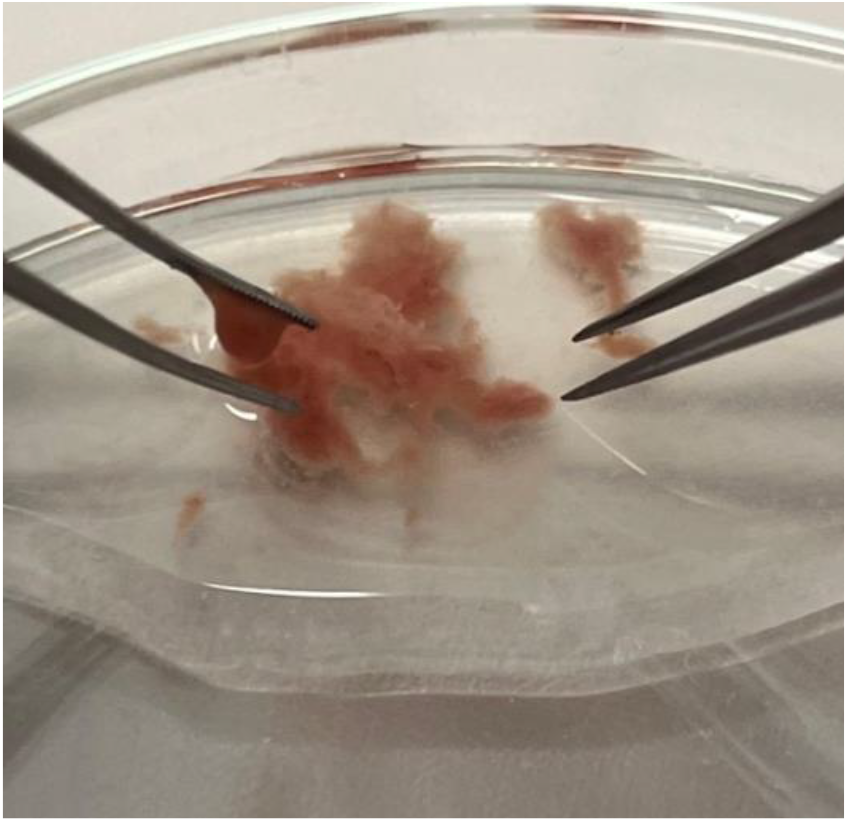
Heart tissue after digestion; tissue readily dissociates with gentle forceps manipulation.

**Figure 5.**
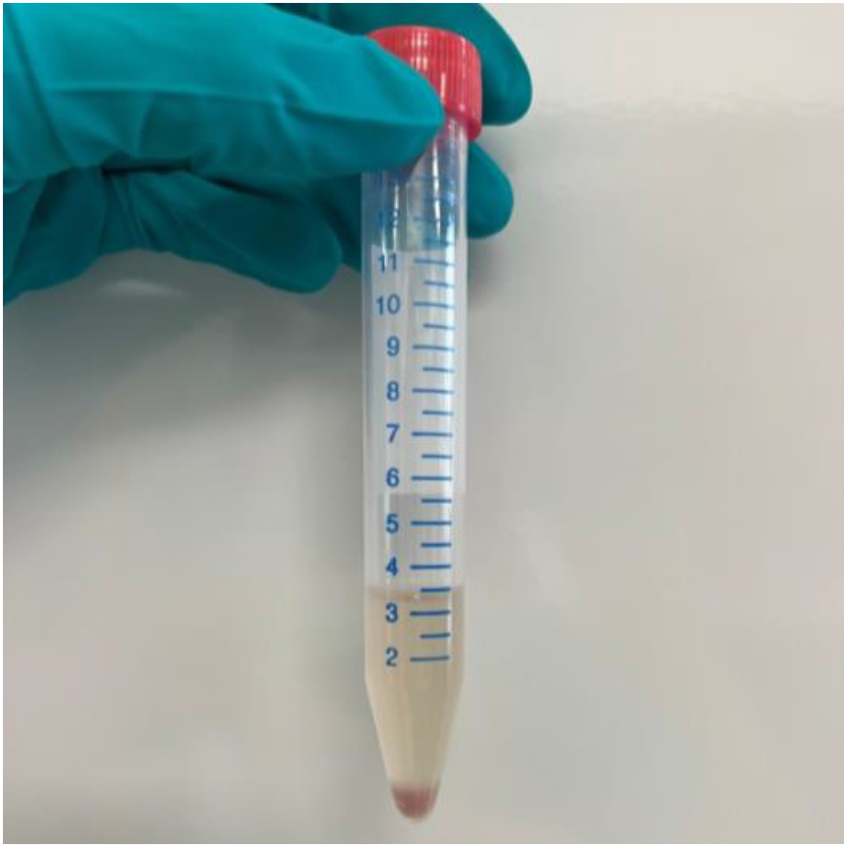
Cell pellet exhibiting distinct pink colour formed after gravity sedimentation. An overly pale or colorless pellet may indicate poor cell viability.

### Gradual Ca^2+^ Reintroduction

Aspirate the supernatant and resuspend the myocyte pellet in 4 mL Ca^2+^-free Tyrode’s solution. Reintroduce Ca^2+^ gradually in a stepwise manner to prevent calcium overload. The final experimental [Ca^2+^] is 1.8 mM.

Increase extracellular Ca^2+^ concentration sequentially in 5–6 steps (4 min incubation at room temperature between steps), for example:

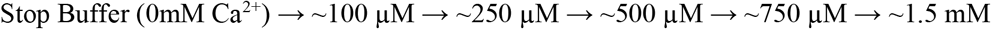

Mix gently after each addition.

Allow cells to settle by gravity, then resuspend in Tyrode’s solution containing 1.8 mM Ca^2+^.

#### Optional

For calcium imaging, load cells with 1–5 µM Fura-2 AM for 20 min (protected from light), followed by transfer to dye-free Tyrode’s solution prior to analysis.

### Anticipated Results

This method typically yields >70% viable rod-shaped ventricular myocytes, with consistent, stimulus-evoked Ca^2+^ transients and preserved responses to β-adrenergic stimulation and caffeine upon reintroduction to physiological [Ca^2+^] (1.8 mM). See Figure 6.

**Figure 6.**
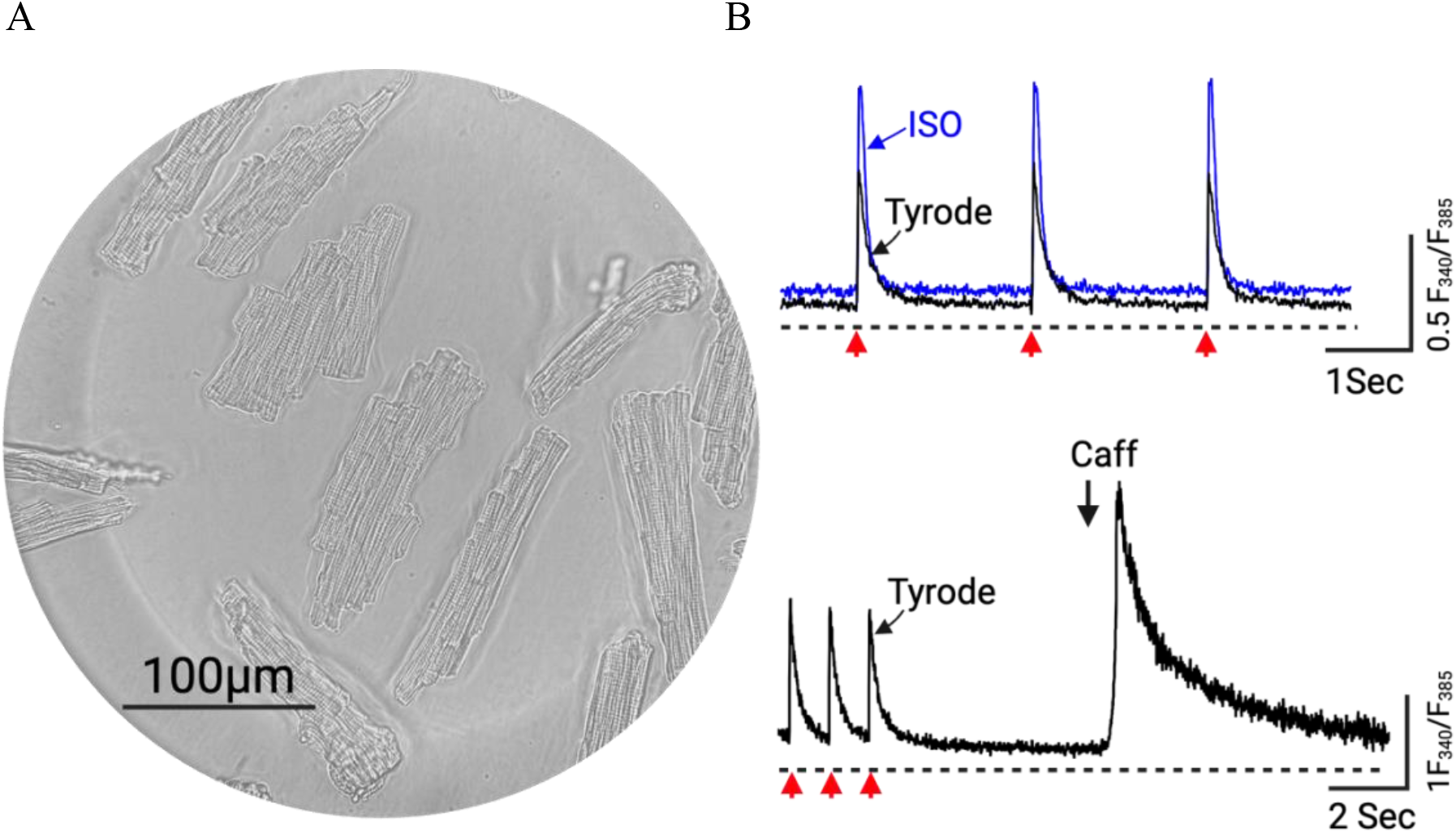
Isolated ventricular myocytes in 1.8mM Ca^2+^ Tryode. A) Representative phase-contrast image of isolated cardiomyocytes. Scale bar = 100 µm. B) Representative traces of Fura-2 loaded cells treated with 50nM isoprenaline, blue line (top) and 10mM caffeine (bottom). Red arrows indicate field stimulation.

### Troubleshooting

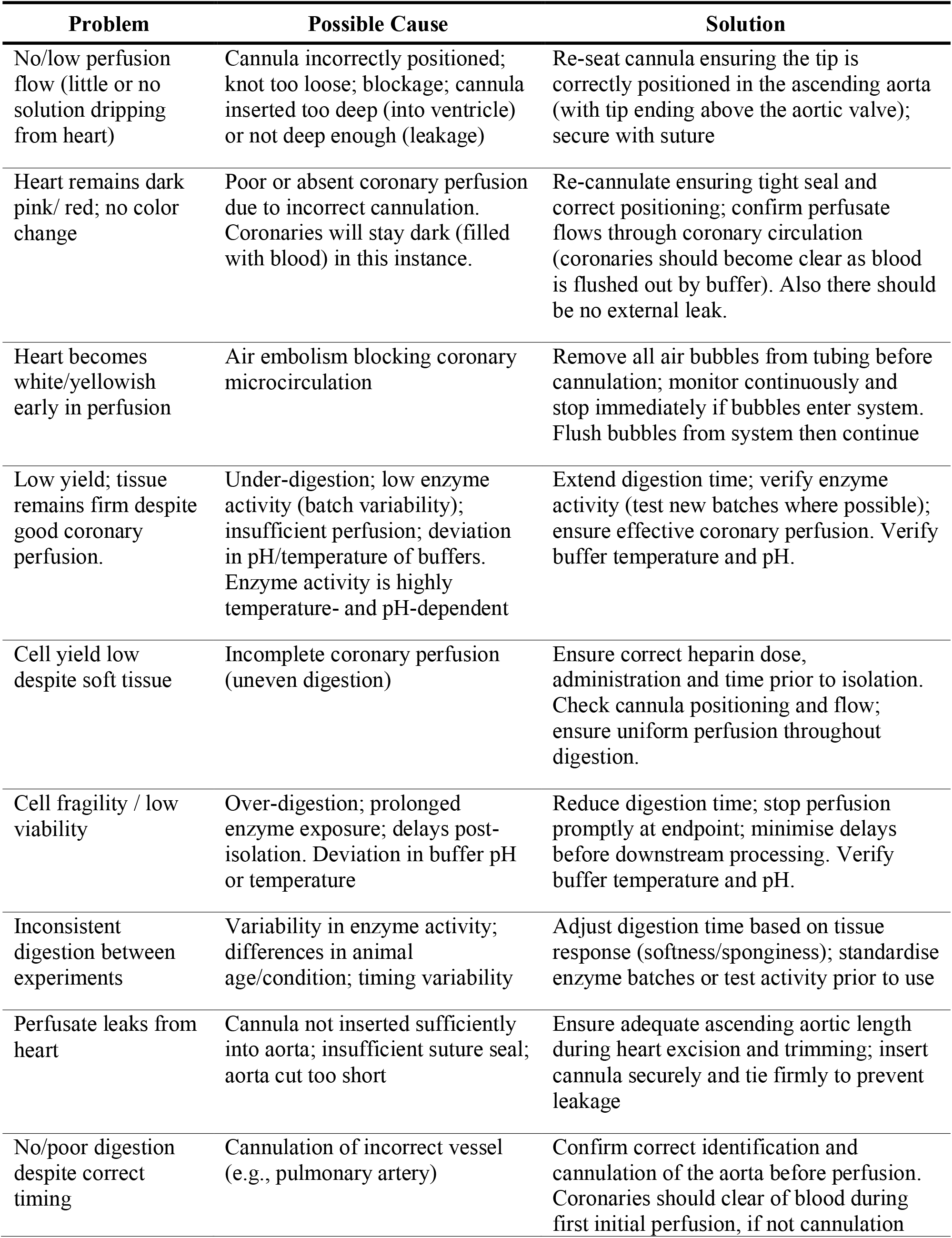
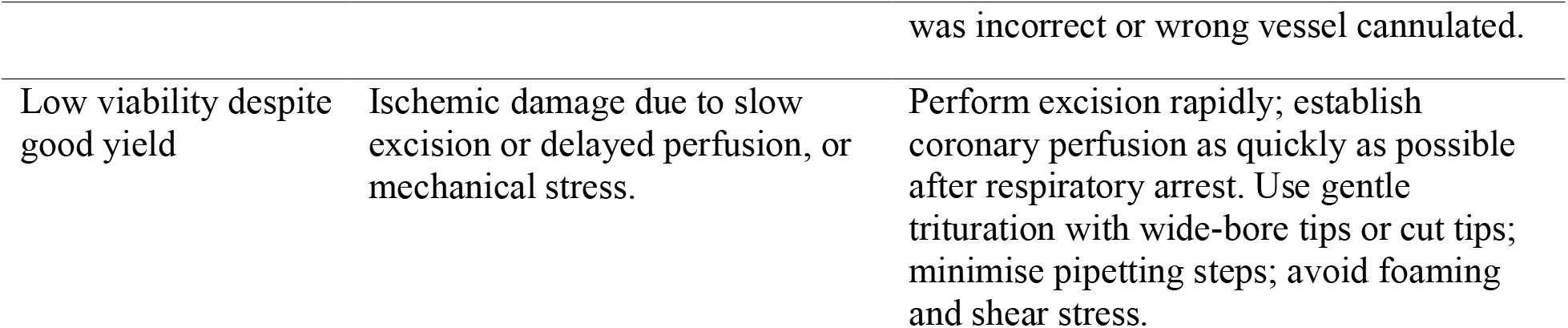

## Notes

### Competing Interest Statement

The authors have declared no competing interest.

